# TBC-2, a Rab GTPase activating protein, regulates the localization of the HLH-30/TFEB and PQM-1 transcription factors in the *C. elegans* intestine

**DOI:** 10.64898/2026.08.14.742015

**Authors:** Soumyendu Saha, Icten Meras, Christian E. Rocheleau

## Abstract

Insulin/IGF signaling (IIS) inhibits the nuclear localization of the DAF-16/FOXO transcription factor to regulate longevity and stress resistance in *C. elegans*. In the intestine, IIS promotes DAF-16 localization to endosomes and loss of TBC-2, a RAB-5 GAP, results in increased endomembrane localization of DAF-16 at the expense of nuclear localization, decreased DAF-16 target gene expression, longevity and stress resistance. Here we found that TBC-2 differentially regulates the localization of the IIS-regulated transcription factors PQM-1 and HLH-30/TFEB. Our results suggest a broader role for TBC-2 in negatively regulating IIS and that TBC-2 likely functions at an upstream point in the IIS pathway.

## Description

Insulin/Insulin-like Growth Factor (IGF) signaling (IIS) is an evolutionarily conserved regulator of metabolism (Barbieri et al., 2003; Murphy & Hu, 2013). In *C. elegans*, the IIS pathway regulates lifespan, fat storage and resistance to various stresses (Friedman & Johnson, 1988; Kenyon et al., 1993; Kimura et al., 1997; Lithgow et al., 1994; Murphy & Hu, 2013; Ogg et al., 1997). The core pathway is defined by the DAF-2 insulin/IGF receptor, the AGE-1 class I PI3 kinase which generates PI(3,4,5)P which in turn recruits PDK-1 and AKT-1/2 kinases to the plasma membrane where PDK-1 phosphorylates and activates AKT-1/2. A major target of *C. elegans* IIS is the inhibition of the DAF-16/FOXO transcription (Kimura et al., 1997; Lin et al., 1997; Morris et al., 1996; Ogg et al., 1997; Paradis et al., 1999; Paradis & Ruvkun, 1998). Phosphorylation of FOXO transcription factors by AKT kinases results in cytoplasmic sequestration (Brunet et al., 1999).

The *C. elegans* intestine, while only 20 cells, is a major metabolic organ and site of many IIS-mediated functions (McGhee, 2007; Murphy & Hu, 2013; Zhang et al., 2022). We previously demonstrated that DAF-16 localizes to endosomes in the intestinal cells (Meras et al., 2022). Interestingly, DAF-16 positive endosomes are not present in every cell or every animal and can vary from just a few to hundreds. We found that TBC-2 regulates DAF-16 localization.

TBC-2 is a RAB-5 GTPase Activating Protein (GAP) that regulates early to late endosome maturation (Chotard et al., 2010). In *tbc-2(tm2241)* deletion mutants, DAF-16 localizes to irregularly shaped endomembranes to a greater extent than wild type and at the expense of nuclear localization (Meras et al., 2022). As such, TBC-2 antagonizes IIS-mediated gene expression, longevity, fat storage and resistance to several stresses (Meras et al., 2022; Traa et al., 2023). To determine if this localization was specific to DAF-16 we looked at two additional transcription factors that are differentially regulated by IIS, PQM-1 and HLH-30.

PQM-1 is a zinc finger transcription factor required for the long lifespan of *daf-2* mutants (Tepper et al., 2013). Localization of PQM-1 is opposite that of DAF-16 whereby IIS promotes nuclear localization of PQM-1 while it antagonizes DAF-16 (Tepper et al., 2013). In *tbc-2(-)* animals, PQM-1::GFP showed some localization to endomembranes (Figure 1A-A’, B). Despite the endomembrane localization of PQM-1, there was a significant increase in the number of animals with nuclear localized PQM-1 and an increase in the nuclear to cytosolic fluorescence intensity ratio of PQM-1 in *tbc-2(-)* mutants as compared to wild type (Figure 1C, D). This corroborates with the antagonistic genetic relation between PQM-1 with DAF-16 localization (Tepper et al., 2013).

**Figure 1.**
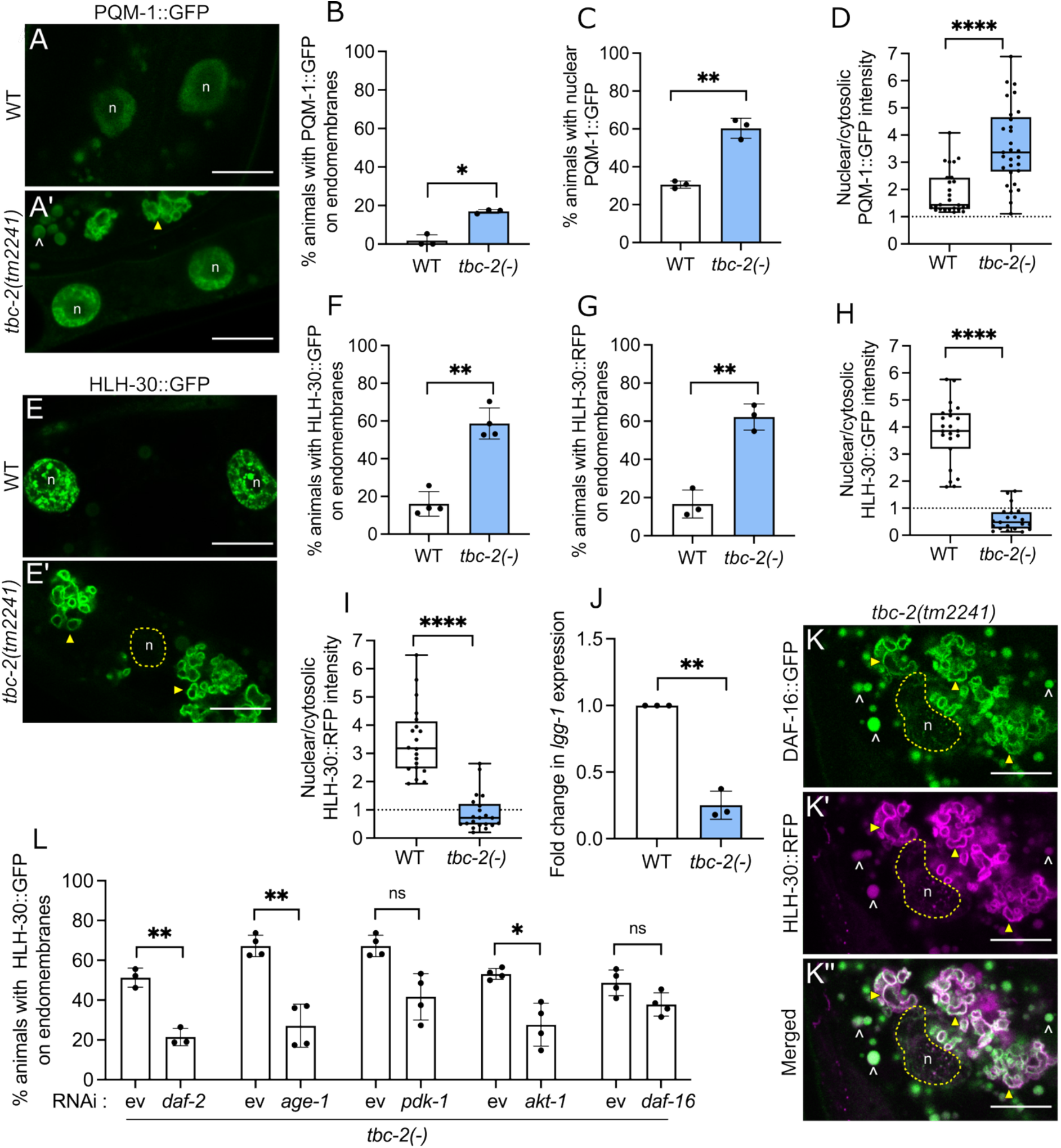
TBC-2 regulates the localization of PQM-1 and HLH-30. (A - A’) Representative confocal images of PQM-1::GFP in the intestine of wild type and *tbc-2(tm2241)* mutants showing nuclear (n) and endomembrane localization (yellow arrow). ^ denotes autofluorescent gut granules. (B, C) Bar graphs showing the percent animals with PQM-1::GFP localization to endomembranes (B) and the nucleus (C). Experimental triplicates with 30-50 animals per replicate. (D) Box and scatter graph representing the ratio of nuclear to cytosolic fluorescence intensity of PQM-1::GFP in wild type (n=26) and *tbc-2(tm2241)* mutants (n =29), each data point is an individual intestinal cell. (E-E’) Representative confocal images of HLH-30::GFP in the intestine of wild type and *tbc-2(tm2241)*. Nuclear (n) and endomembrane localization (yellow arrows). (F, G) Bar graphs showing the percent animals with endomembrane localization of HLH-30::GFP (F), and HLH-30::RFP (G) in wild type and *tbc-2(tm2241)* mutants. Experimental replicates with 30-60 animals per replicate. (H, I) Box and scatter graphs representing the ratio of nuclear to cytosolic fluorescence intensity of HLH-30::GFP (H) and HLH-30::RFP (I) in wild type and *tbc-2(tm2241)* mutants, n > 20 for each condition. (J) Bar graph showing the fold change in *lgg-1* mRNA expression levels between wild type and *tbc-2(tm2241)* mutant, in 3 biological replicates. (K-K”) Representative confocal images DAF-16::GFP and HLH-30::RFP colocalization in *tbc-2(tm2241)* as shown by the yellow arrow. ^ marks the auto fluorescent gut granules and n denotes nuclei. (L) Bar graph showing the percent animals with endosomal HLH-30::GFP in the *tbc-2(tm2241)* background, fed control (ev, empty vector) RNAi or RNAi targeting components of the IIS pathway. Each replicate represents >25 worms per condition. Paired Student T-test was used to analyse data in (B, C, F, G, J, and L) and unpaired Student T-test was used to analyse data in (D, H, and I). Scale bar, 10 µm. *P < 0.05, **P<0.01 and ****P<0.0001.

HLH-30 or TFEB (Transcription factor EB) is a transcription factor involved in lysosomal biogenesis and acts in a combinatorial manner with DAF-16 in response to the starvation stress (Lapierre et al., 2013; Lin et al., 2018; O’Rourke & Ruvkun, 2013). In wild type we observed HLH-30::GFP was mostly nuclear with some cytoplasmic and endomembrane localization. In *tbc-2(-)* mutants, using both GFP and RFP tagged HLH-30 we observed an increase in the number of animals with HLH-30 localized to endomembranes (Figure 1E-E’, F, G). Like DAF-16, HLH-30 showed reduced nuclear to cytosolic fluorescence intensity in *tbc-2* mutant animals (Figure 1H,I). To test if the reduced nuclear localization of HLH-30 affected downstream HLH-30 dependent gene expression, we analysed the mRNA expression of *lgg-1*. LGG-1 is ATG8 in yeast and LC3 in mammals and is responsible for autophagosome formation (Melendez et al., 2003). LGG-1 expression is dependent on HLH-30, such that *lgg-1* mRNA levels are suppressed in *hlh-30(RNAi)* (Lapierre et al., 2013).

There was a significant decrease in the *lgg-1* mRNA levels in *tbc-2(tm2241)* mutants consistent with the reduced nuclear localization of HLH-30 (Figure 1J).

We observed both DAF-16 and HLH-30 co-localized on endomembranes in *tbc-2(-)* mutants indicating that TBC-2 might regulate these transcription factors by a common underlying mechanism (Figure 1 K-K”). To test if IIS regulates HLH-30 localization on endosomes as seen with DAF-16 (Meras et al., 2022), we use RNAi to systematically knock down components of the IIS pathway-DAF-2, AGE-1, PDK-1, AKT-1 and DAF-16 in *tbc-2(-);* HLH-30::GFP animals. Compared to empty vector control, RNAi mediated knock-down of *daf-2, age-1* and *akt-1* significantly decreased the number of *tbc-2(-)* mutant animals with HLH-30 on endomembranes (Figure 1L). *pdk-1(RNAi)* significantly suppressed HLH-30 endomembrane localization in three of four experiments (Figure 1L). *daf-16* RNAi did not decrease the endomembrane localization of HLH-30 (Figure 1J). Thus, IIS signaling promotes the localization of HLH-30 to endomembrane compartments as it does with DAF-16.

In conclusion, this study builds upon our previous findings that TBC-2 regulates DAF-16 localization to endomembranes (Meras et al., 2022). We demonstrated that TBC-2 regulates the localization of two additional IIS regulated transcription factors, HLH-30 and PQM-1, further supporting a role for TBC-2 in antagonizing IIS in the intestine. Similar to DAF-16, we found that HLH-30 localizes to uncharacterized endomembrane compartments in *tbc-2* mutants at the expense of its nuclear localization. The reduced expression of the HLH-30 target, *lgg-1*, in *tbc-2* mutants suggests that this impacts HLH-30 transcriptional activity.

We found that HLH-30 colocalizes with DAF-16 on the same endomembranes in *tbc-2* mutants suggesting a similar mode of regulation. HLH-30/TFEB and DAF-16/FOXO are inhibited by phosphorylation and binding with 14-3-3 proteins that sequester them in the cytoplasm (Brunet et al., 1999; Colino-Lage et al., 2024; Li et al., 2007; Lin et al., 2018; Martina et al., 2012; Roczniak-Ferguson et al., 2012). We previously demonstrated that FTT-2/14-3-3 and DAF-16 phosphorylation regulate DAF-16 localization to endomembranes (Meras et al., 2022). We anticipate that HLH-30 is also sequestered in a similar manner.

However, we don’t know whether 14-3-3 mediates HLH-30 and DAF-16 binding to these endomembrane structures or if there are additional factors involved.

DAF-16 and HLH-30 forms a complex and regulates overlapping stress response genes (Lin et al., 2018). Whether they form a complex on endomembranes remains unclear. RNAi of *daf-16* did not affect the localization of HLH-30 to endomembranes suggesting that there may not be a codependence for endomembrane localization. However, this should be further tested with null alleles.

Like DAF-16, we also found that IIS partially regulates the localization of HLH-30 to endomembranes. Thus, it is possible that other metabolic or stress response pathways such as AMPK, Tor, or JNK could be involved (El-Houjeiri et al., 2019; Greer et al., 2007; Lapierre et al., 2013; Martina et al., 2012; Oh et al., 2005; Roczniak-Ferguson et al., 2012; Settembre et al., 2012). Unlike DAF-16 and HLH-30, IIS promotes nuclear localization of PQM-1 (Tepper et al., 2013). Although we detected some endomembrane localization of PQM-1 in *tbc-2* mutants, we found that the nuclear localization of PQM-1 was increased. This is consistent with our previous finding that *tbc-2* mutants had increased expression of *vha-6*, a top PQM-1 target (Meras et al., 2022; Tepper et al., 2013). Taken together these results suggest that TBC-2 antagonizes IIS at a common point upstream of the DAF-16, HLH-30 and PQM-1 transcription factors.

## Methods

Worms were maintained at 20°C on NGM plates as described in WormBook (www.wormbook.org). The strains were fed HB101 *E. coli* as a food source.

For RNAi feeding experiments fourth larval stage (L4) worms were placed on RNAi plates and transferred to fresh plates each day for 3 days. L4 progeny from the day 3 plates were scored for the experiment. RNAi feeding clones were obtained from the Ahringer RNAi library (Kamath et al., 2001). *pdk-1* RNAi clone (pQcSS01) was generated by PCR of worm lysates using primers SSA38 5’-CAACTAAGATCTATTTCGTGATCGGACTTGTTG-3’ and SSA395’-GTCATACTCGAGGCTAGCATCATTTCCCAAATTCATC-3’ and ligated to the L4440 vector from Addgene (https://www.addgene.org/1654/).

Live imaging of the L4 hermaphrodite worms was done at room temperature. The worms were transferred to a 2% agarose pad and immobilized with 10 mM levamisole. They were scored for nuclear or endosome localization of PQM-1, DAF-16 and HLH-30 under the fluorescent ZeissA1 axio Imager. To measure the fluorescence intensity of nuclear PQM-1 and HLH-30 L4 worms were imaged using a Zeiss Axio Observer Z1 LSM780 laser scanning confocal microscope in the Molecular Imaging Platform of the RI-MUHC. Image analysis was performed using Fiji (ImageJ).

For qualitative RT PCR, RNA was extracted from young adult *C. elegans* using TRIZOL as explained in (Meras et al., 2022). LGG-1 was amplified using forward primer 5’-ACCCAGACCGTATTCCAGTG-3’ and reverse primer 5’-ACGAAGTTGGATGCGTTTTC-3’.

## Reagents

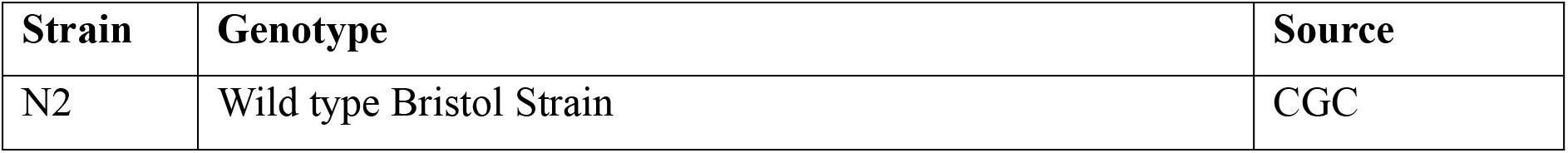
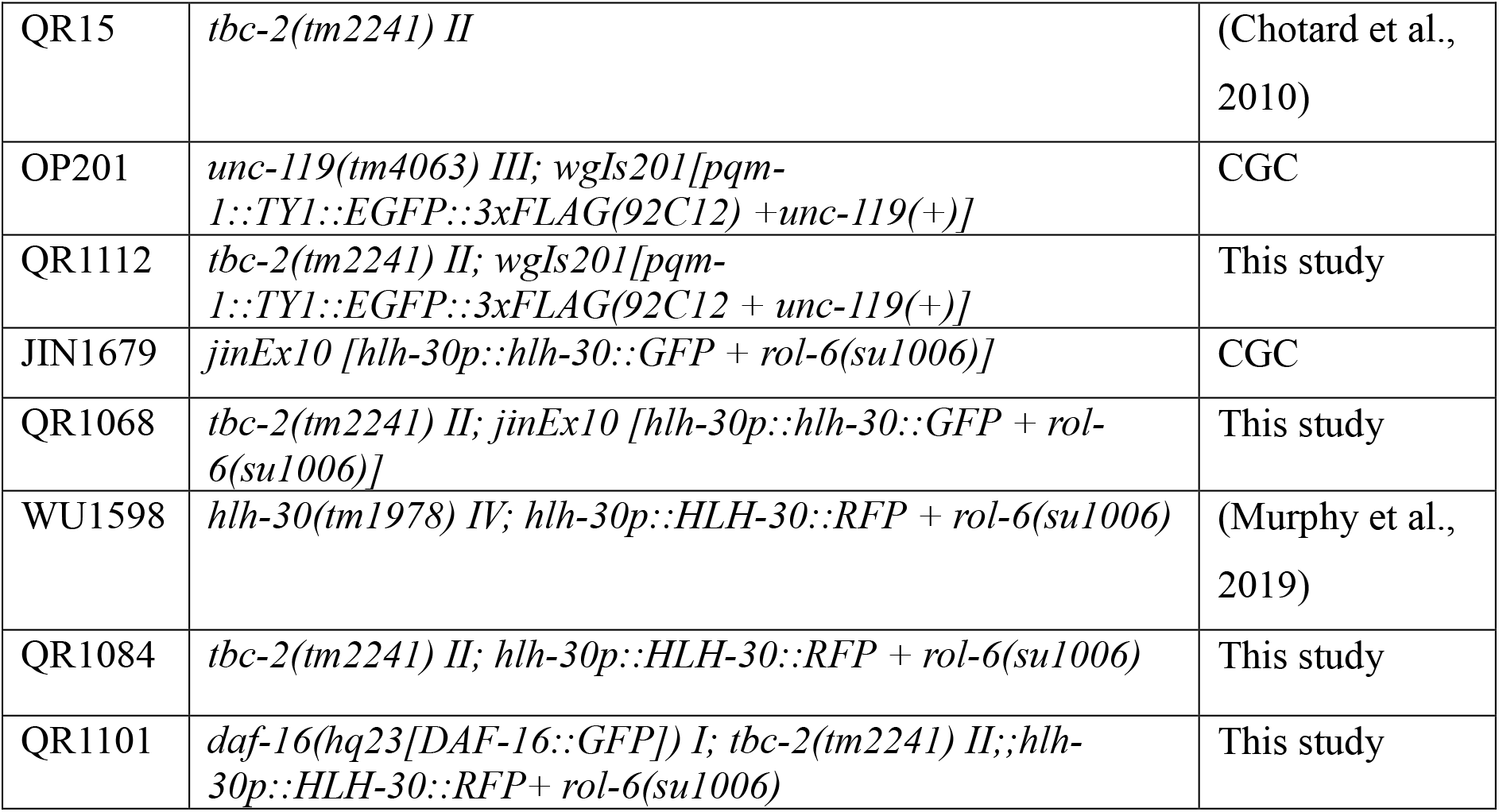

| RNAi clone | Gene | Source |
| --- | --- | --- |
| L4440 | Empty vector control | (Timmons & Fire, 1998) |
| III-7G12 | *daf-2* | (Kamath et al., 2003) |
| II-7J02 | *age-1* | (Kamath et al., 2003) |
| V-7I17 | *akt-1* | (Kamath et al., 2003) |
| I-5M24 | *daf-16* | (Fraser et al., 2000) |
| pQcSS01 | *pdk-1* | This study |

## Acknowledgements

We would like to thank Jung Hwa Seo for technical assistance. We thank Kerry Kornfeld and Abhinav Diwan (Washington University) and Jeremy Van Raamsdonk (McGill University) for sharing strains and RNAi clones. Some strains were provided by the Caenorhabditis Genetics Center (CGC), which is funded by NIH Office of Research Infrastructure Programs (P40 OD010440).

